# Efficient 3ʹ-end tailing of RNA with modified adenosine for nanopore direct total RNA sequencing

**DOI:** 10.1101/2024.02.24.581884

**Authors:** Yinan Yuan, Reed Arneson, Emma Burke, Alexander Apostle

## Abstract

Direct sequencing of total cellular RNA enables a better understanding of a broad spectrum of RNA species controlling cellular processes and organismal function. Current nanopore direct RNA sequencing method, however, only captures polyadenylated RNA for sequencing. To address this issue, we developed a unique 3’-end RNA tailing method to capture total RNA for nanopore direct RNA sequencing. Due to the distinct electrical signature of the added tail on nanopore, this method allows simultaneous detection of both non-polyadenylated and polyadenylated RNAs. We demonstrated the effectiveness of this method in capturing the dynamics of transcription and polyadenylation of chloroplast RNAs in plant cell. With its high efficiency in retaining total RNA on nanopore, this method has the potential to be broadly applied to RNA metabolism and functional genomics studies.

---

Transcriptional and post-transcriptional processes in cell are complex [1-3], generating total cellular RNA species that are critical regulators of cell fate and organismal development [4-10]. Tracking the dynamics of the total transcriptome and RNA metabolism is crucial for understanding cell biology and organismal complexity. The success of this endeavor relies mostly on our ability to faithfully capture the sequence, structure and chemical diversity of total cellular RNA for analysis.

Nanopore direct RNA sequencing which sequences intact RNA strands and provides both nucleotide sequence and base modifications at the isoform level [11, 12], is the least biased RNA sequencing platform for studying total cellular RNA dynamics. The total RNA in cell consists of protein coding messenger RNAs (mRNAs), generally polyadenylated at the 3’ end, and various forms of noncoding RNAs (ncRNAs), which are largely non-polyadenylated at the 3’ end. However, the current direct RNA sequencing kit from Oxford Nanopore Technologies (ONT) only captures polyadenylated (poly(A)) RNAs, and most ncRNAs, or any mRNAs without poly(A) tails (poly(A)^-^) are excluded for sequencing. Technically, in order for poly(A)^-^ RNA molecules to be sequenced on nanopore, either custom adaptors designed for specific groups of RNAs [13, 14], or custom polynucleotide tails synthesized through special enzymes, have to be added to the 3’ end of RNA [15-17].

RNA nucleotidyl transferases such as poly(A) polymerases add ribonucleotides to the 3’ end of RNA in a template-independent manner, and have been routinely used for in vitro 3’ end tailing and labelling [18-20]. Recently, poly(A) tailing has been used to capture poly(A)^-^ RNA for nanopore direct RNA sequencing [17, 21]. With poly(A) tailing, however, the native polyadenylation of poly(A) RNA at 3’ end is lost. Thus, to preserve the native polyadenylation of poly(A) RNA species in total cellar RNA population, adding polymers of modified or unnatural nucleotides to the 3’ end of RNA is required [15, 16]. The existing non-poly(A) tailing methods, however, are suboptimal in efficiency of tailing poly(A)^-^ RNA and in sequencing throughput on nanopore [15, 17]. Taken together, the technology for direct sequencing of total cellular RNA on nanopore still awaits improvement.

To address the technical challenges in direct nanopore total RNA sequencing, we focused on using yeast poly(A) polymerase (YPAP) to add modified ATP analogs efficiently to the 3’ end of RNA, in particular poly(A)^-^ RNA, with the goal to integrate our method with the current ONT direct RNA kit for simultaneous detection of both poly(A)^-^ and poly(A) RNAs. Here we present the unique activity of yeast poly(A) polymerase in synthesizing a short tail of 2′-O-methyladenosine (referred as poly(mA) hereafter) to the 3’ end of RNA. We demonstrated the high efficiency of our method in tailing poly(A)^-^ RNA, the distinct electrical signature of poly(mA) tail on nanopore, and the effectiveness of poly(mA) tailing in capturing the total RNA population and in-depth profiling of organellar RNAs.

## RESULTS

### YPAP adds poly(mA) tail to the 3’ end of poly(A)^-^ RNA with high efficiency

We tested the ability of YPAP in adding modified ATPs to the 3’ end of RNA, especially poly(A)^-^ RNA. The modified ATPs surveyed in our study included 2′-OMe-ATP (2′-OMeA), N6-methyl-ATP (m6A) and 2-aminoadenosine-5’-triphosphate (2-amino-ATP). These ATP analogs are structurally distinct from ATP (Supplementary Fig. S1), and some of these nucleotides have been confirmed to generate distinguished current traces on nanopore [13, 22].

We first assessed the activity of YPAP in catalyzing a 20-mer non-polyadenylated RNA oligo, alongside the poly(A) polymerase from *E. coli* (EPAP). Under the same experimental conditions, the ability of YPAP in catalyzing same amount of RNA oligo, varied with ATP analogs used, compared with EPAP. Overall, YPAP was much less efficient than EPAP in using ATP, 2-amino-ATP and m6A, but was highly efficient in adding poly(mA) tail (Fig. 1a). We observed that the substrates were efficiently tailed with poly(mA) in 5-minute of reaction; while much of the oligo substrates remained uncatalyzed with other two analogs (Fig. 1a). In addition, the poly(mA) tails added by YPAP were generally longer than those added by EPAP.

**Figure 1.**
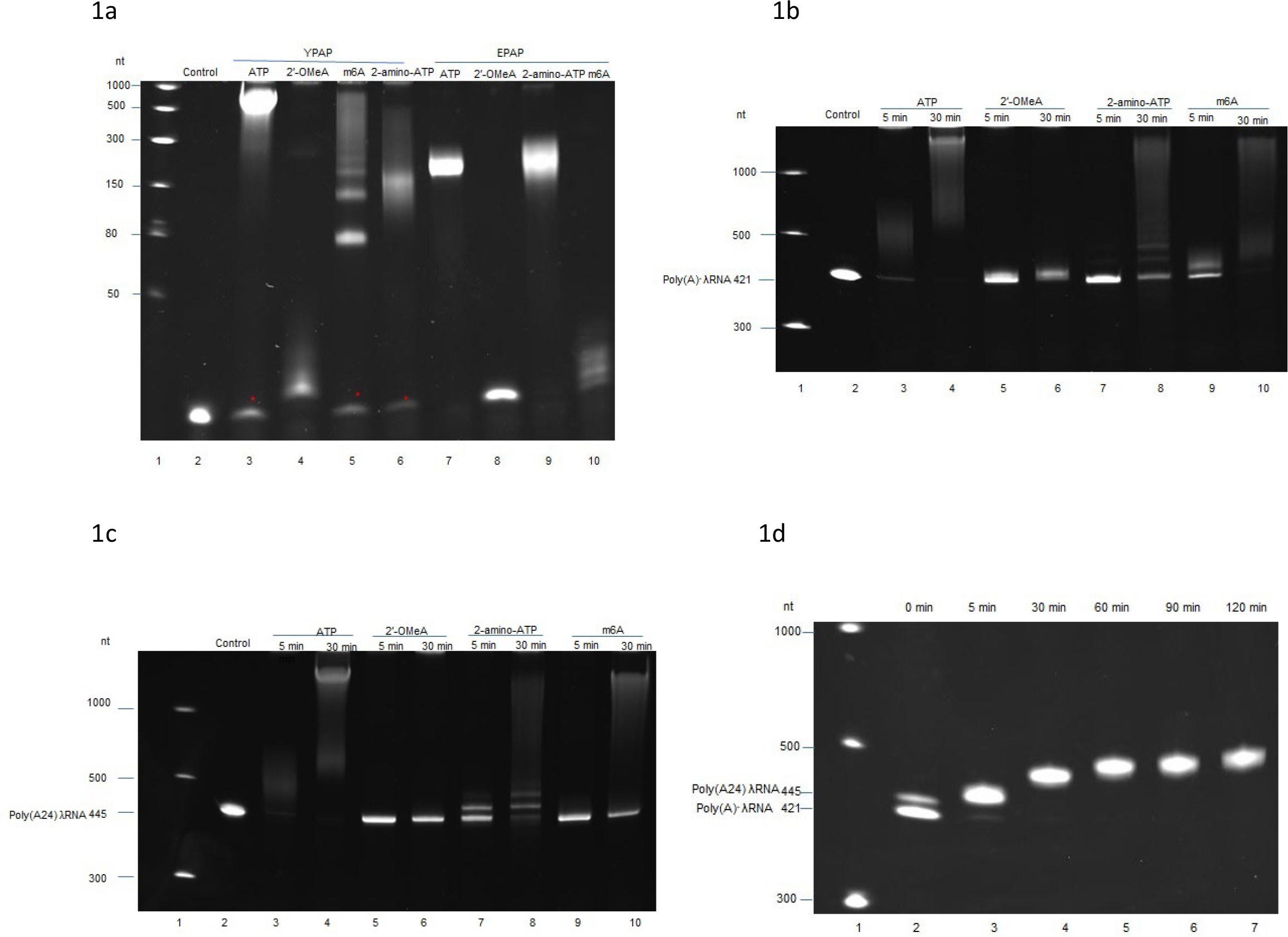
Characterizing the enzyme activity of YPAP in adding 2’-OMeA to the 3’ end of RNA. **(a)** The nucleotide-adding ability of YPAP vs. EPAP in catalyzing the 20-mer RNA oligo. Red asterisks indicate uncatalyzed substrate of 20-mer RNA identified in tailing reaction of ATP, 2-amino-ATP or m6A by YPAP after 5 min of reaction at 37°C. **(b)** The efficiency of YPAP in catalyzing the poly(A)^-^ λRNA substrate using 2’-OMeA, alongside ATP, 2-amino-ATP or m6A, was evaluated at 5 min and 30 min of incubation at 37°C. **(c)** The ability of YPAP to add 2’-OMeA to the 3’ end of poly(A24) λRNA was examined side-by-side with ATP, 2-amino-ATP or m6A, over a 5 min and 30 min of reaction at 37°C. **(d)** YPAP showed distributivity and processivity in adding 2’-OMeA to both poly(A)^-^ and poly(A) RNA of the substrate mixture containing poly(A)^-^ λRNA and poly(A24) λRNA at a ratio of 2:1 over a time series of tailing reaction at 37°C.

Next, we tested the efficiency of YPAP in catalyzing the addition of ATP analogs to longer non-polyadenylated RNAs. We used in vitro transcribed λRNAs as substrates. The pattern of analog addition by YPAP to the long RNA substrate was similar to that observed with RNA oligos. YPAP efficiently added poly(mA) tail to poly(A)^-^ RNAs but much less efficiently used 2-amino-ATP and m6A (Fig. 1b). Despite the relatively short length of the poly(mA) tail added to the longer RNA, the shift in the substrate RNA band on a 6% PAGE gel was perceptible following the 30-minute reaction, suggesting that an increasing number of 2′-OMeA were added distributively to the substrates as the tailing reaction progressed (Fig. 1b).

### YPAP adds 2′-OMeA to poly(A)^-^ RNAs more favorably than to poly(A) RNAs

To examine the efficiency of YPAP in adding poly(mA) tail to poly(A) RNAs, we used in vitro transcribed polyadenylated λRNA (poly(A24) λRNA) as the substrate. Based on gel analysis of tailing reactions, the poly(mA) tail added to poly(A24) λRNA by YPAP was shorter than that added to poly(A)^-^ RNA. We found that under the same reaction time, the poly(A24) RNA band on the gel migrated only slightly after 30 minutes of incubation (Fig. 1c). It seems the presence of poly(A) on the 3’ end of RNA affects the incorporation rate of YPAP for some unknown reasons.

To understand how YPAP uses 2′-OMeA when catalyzing natural RNA populations consisting of both poly(A) and poly(A)^-^ RNAs, we performed time-series of tailing experiment using a substrate mixture of poly(A)^-^ λRNA and poly(A24) λRNA (2:1 ratio). As shown in Fig. 1d, the poly(A)^-^ RNA band diminished faster than the poly(A24) RNA band after 5 minutes of reaction time, which implies that poly(mA) tailing of poly(A)^-^ substrates by YPAP was favored initially. Over the time series, the poly(A)^-^ RNA became gradually undetectable on the gel and the poly(A) RNA shifted noticeably upwards, meaning that both poly(A) and poly(A)^-^ RNA were progressively catalyzed by YPAP using 2′-OMeA. Interestingly, we have consistently observed that, instead of adding long polymers of ATP or its analogs, YPAP generally added a narrow range of poly(mA) tails to RNA molecules. Even after a 2-hour incubation at 37°C, YPAP produced controlled poly(mA) tails on both poly(A)^-^ and poly(A) RNAs in a narrow range (roughly 50 to 70 nt), without any high-molecular-weight tailed products detected (Fig. 1d). This finding is noteworthy since long polynucleotide tails have been linked to low sequencing throughput on nanopore.

As a comparison, we inspected the efficiency of EPAP in adding poly(mA) tails to the same amount of substrate mixture, as well as the efficiency of YPAP and poly(U) polymerase (PUP) in synthesizing poly(I) tails to the 20-mer RNA substrate and to substrate mixtures of poly(A)^-^ and poly(A) λRNAs (Supplementary Fig. S2). In every case that we examined, poly(mA) tailing by YPAP outperformed other non-poly(A) tailing in efficiency of catalyzing poly(A)^-^ RNA (Supplementary Fig. S2a-d). Notably, despite its slow incorporation rate, YPAP is capable of poly(mA) tailing poly(A) RNA through increased reaction time, with a comparable performance to the poly(A) tailing by EPAP, although the latter catalyzes the substrate at a much higher rate (Supplementary Fig. S2e).

### The ionic current traces of the poly(mA) tail on nanopore can be distinguished from the poly(A) signal

To capture and retain the native information of total cellular RNAs, the poly(mA) tail added to the 3’ end of RNA must be distinguishable from the native poly(A) tail on nanopore. It has been demonstrated that nanopore is sensitive to nucleotide modifications including 2′-OMeA modification [13, 21, 22]. Therefore, we reasoned that the poly(mA) would generate current traces distinct from the native poly(A) signal on nanopore. To test this hypothesis, we constructed three types of in vitro transcribed λRNA libraries including poly(A)^-^ λRNA tailed with either poly(mA) or poly(A), and poly(A)-tailed λRNA (poly(A) RNA) further tailed with poly(mA). The poly(mA)-tailed RNAs were fully integrable with ONT direct RNA library preparation (Fig. 2a), and the resulting libraries were run on a MinION sequencer.

**Figure 2.**
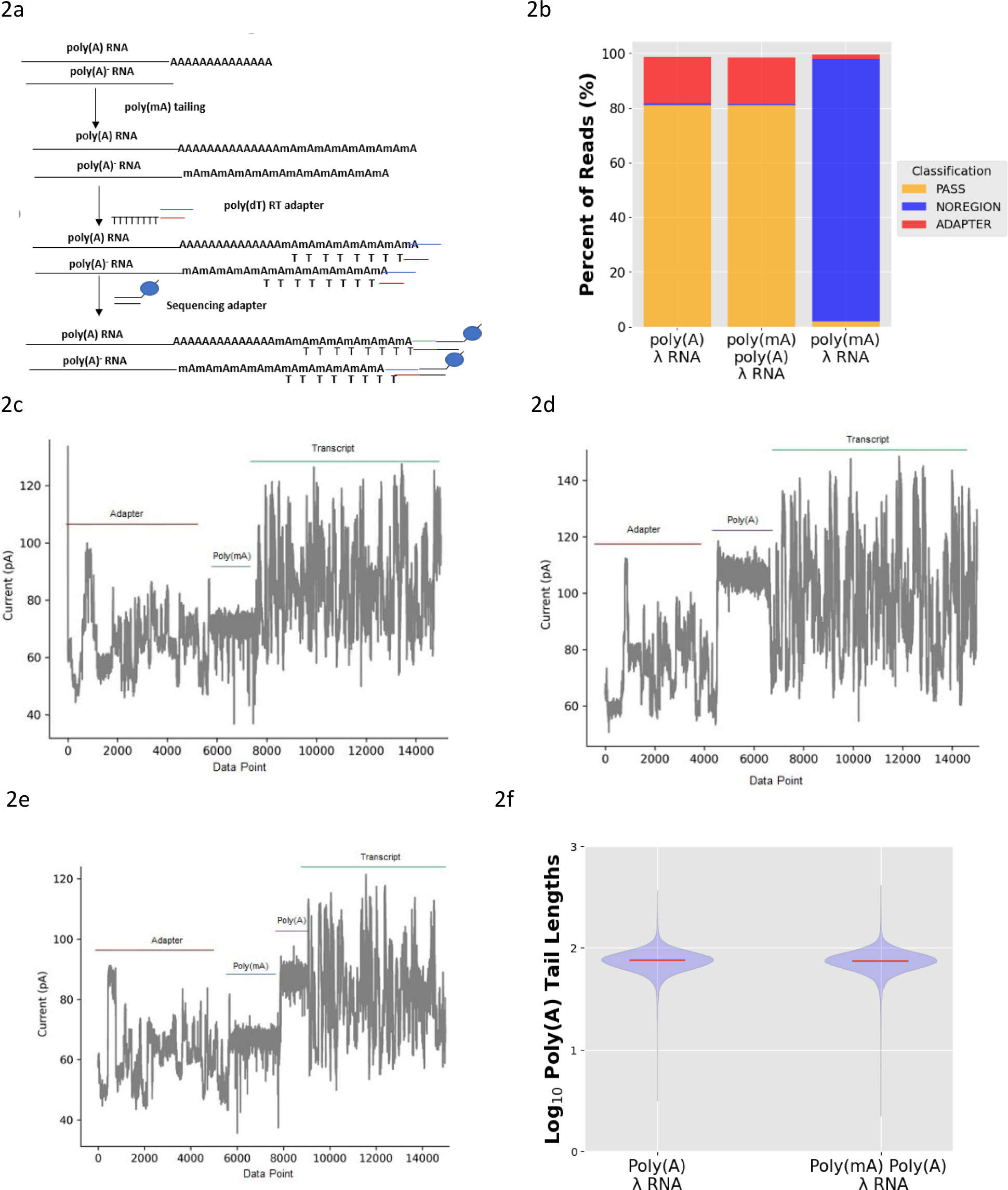
Direct poly(mA)-tailed RNA sequencing on nanopore. **(a)** Library preparation for poly(mA)-tailed RNAs. **(b)** Classification of λRNA reads from three tailed λRNA libraries employing nanopolish. PASS: poly(A) RNA, NOREGION: poly(A)^-^ RNA. **(c)** Nanopore ionic current traces of a poly(mA)-tailed λRNA with the specific poly(mA) signal identified. **(d)** Nanopore ionic current traces of a poly(A)-tailed λRNA with the classical poly(A) tail signal detected. **(e)** Nanopore Ionic current traces of a poly(mA) tail detected between the adapter and poly(A) region of a poly(mA)-tailed poly(A) λRNA. **(f)** The violin plot shows the distribution of the poly(A) tail length between a poly(A) λRNA library and a poly(mA)-tailed poly(A) λRNA library. The mean tail length is specified by the red line.

The electrical signal produced by a poly(mA) tail on nanopore could be distinguished from a poly(A) signal by nanopolish, a signal-level analysis tool which classifies reads containing a poly(A) tail into the “PASS” category and estimates the length of that poly(A) tail based on segmenting signals [23]. We found that as many as 93% of reads from the poly(mA)-tailed λRNA library were classified as “NOREGION” (without a poly(A) signal), while at least 81% of reads from the poly(A)-tailed λRNA library were recognized as “PASS” (containing a poly(A) tail) (Fig. 2b). This result suggests that the current traces of the poly(mA) tail added to poly(A)^-^RNA were so distinct that nanopolish was able to differentiate it from the poly(A) signal. Inspecting the raw electrical signal of the 3’ end of sequenced RNA tailed either with poly(mA) or poly(A) supports the finding that the poly(mA) signal on nanopore is specific and distinguishable, with the mean ionic current usually below 80 pA, much lower than the mean current traces of poly(A) tail (Fig. 2c-d).

We also investigated whether the signal of the poly(mA) added to a poly(A) RNA interfered with poly(A) segmentation and tail length estimation by nanopolish. Interestingly, we found that a significant number of reads (81%) from a poly(mA)-tailed poly(A) λRNA library were still recognized as poly(A) containing reads. This is approximately the same percentage of “PASS” reads that we identified in the control poly(A) λRNA library (Fig. 2b). We also examined signal variations between the adapter and poly(A) tail and identified the characteristic poly(mA) current traces distinct from the 100 pA, mean current traces of a neighboring poly(A) segment (Fig. 2e). In most poly(mA)-tailed poly(A) RNA molecules, the segment of poly(mA) was rather narrow, probably due to the much shorter poly(mA) tail added to poly(A) RNA observed in earlier tailing experiment. We further compared the poly(A) tail length distribution between the poly(A) λRNA library and the poly(mA)-tailed poly(A) λRNA library, and determined that poly(A) tail length was distributed similarly between the two libraries (Fig. 2f), confirming the proper poly(A) segmentation of poly(mA)-tailed poly(A) RNAs by nanopolish.

Taken together, we conclude that the distinct signal generated by the poly(mA) tail on nanopore can be differentiated and properly segmented from the current traces of poly(A). As a result, this allows both poly(A) and poly(A)^-^ RNAs to be detected on nanopore simultaneously and the length and pattern of polyadenylation of poly(A) RNAs to be conveniently analyzed by signal level computational tools.

### Poly(mA) tailing captures the depth of the cellular RNA population

To test the effectiveness of poly(mA) tailing in capturing the total RNA population in natural cellular content, we performed poly(mA) tailing of three types of cellular RNA samples including mRNAs, total RNAs, and total RNAs with ribosomal RNA depleted (rRNA^-^). We constructed and sequenced tailed RNA libraries along with two control libraries (Table 1). All RNA samples were derived from the same *Populus* leaf tissue. Sequencing run statistics and genome mapping information for each library are summarized in Supplementary Table S1. For each library, we assigned sequenced reads as either poly(A) RNA or poly(A)^-^ RNA based on nanopolish classification, and this could be further confirmed through raw signal analysis of the poly(mA) and the presence/absence of a native poly(A) region. Some gene loci such as Potri.010G150500, a stress responsive A/B barrel domain protein, transcribed both poly(A) and poly(A)^-^ RNA isoforms which were captured effectively by poly(mA) tailing (Fig. 3a, 3b).

**Table 1.** Natural cellular RNA populations captured by poly(mA) tailing on nanopore.

| Library type | RNA | Poly(mA) tailing | poly(dT) RTA adapter | RNA to be captured |
| --- | --- | --- | --- | --- |
| Control-mRNA | Enriched mRNA | - | + | only poly(A) RNA |
| Tailed-mRNA | Enriched mRNA | + | + | poly(A) RNA, and poly(A)<sup>-</sup> RNA |
| Tailed-total RNA-rRNA<sup>-</sup> | Total RNA (rRNA<sup>-</sup>) | + | + | poly(A) RNA and poly(A)<sup>-</sup> RNA |
| Tailed-total RNA | Total RNA | + | + | poly(A) and poly(A)<sup>-</sup> RNA |
| Control-total RNA | Total RNA | - | + | only poly(A) RNA |

**Figure 3.**
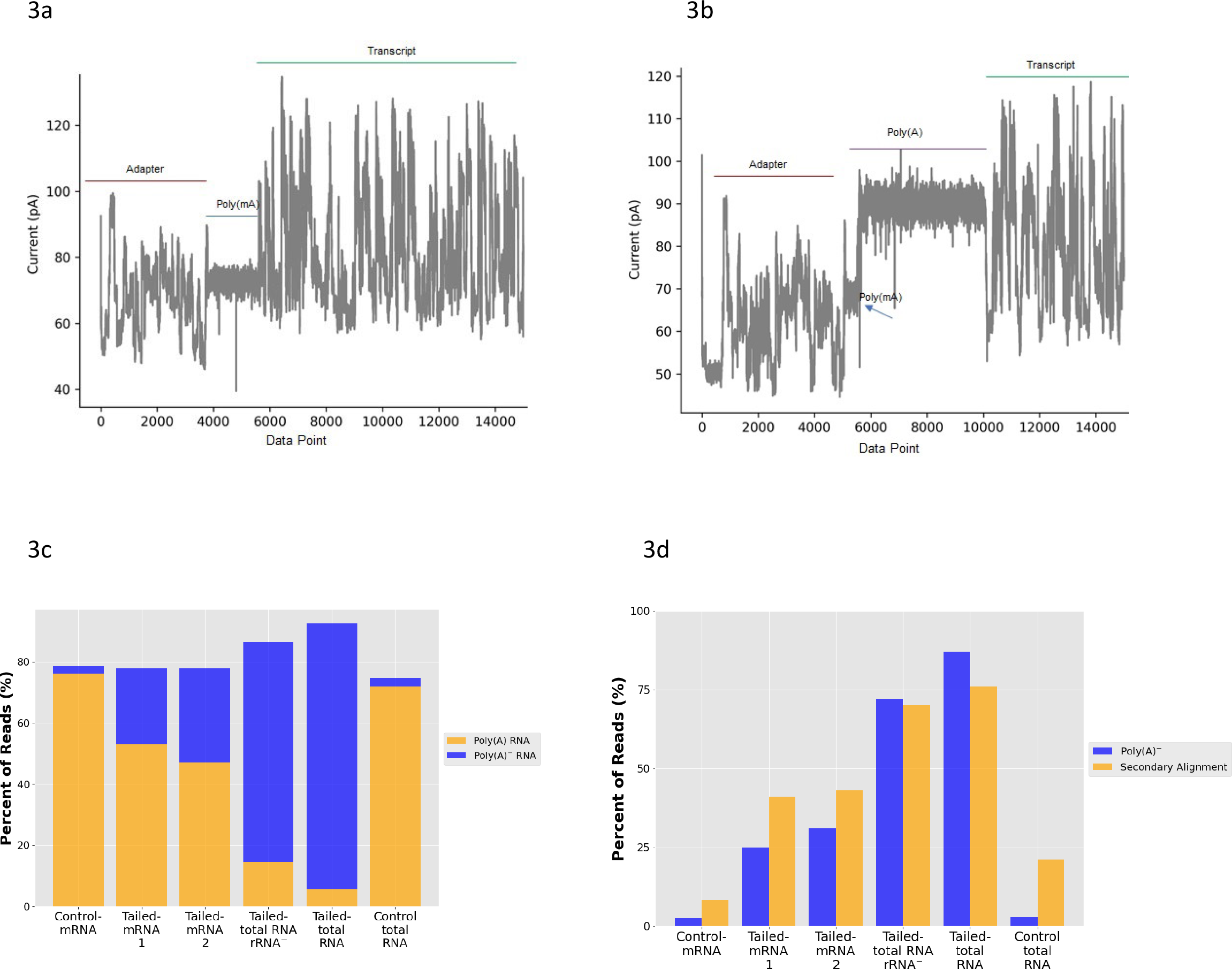
Poly(mA) tailing captures cellular RNAs on nanopore. **(a)** Raw electrical signal of poly(mA) identified from one poly(mA)-tailed poly(A)^-^ RNA isoform of Potri.010G150500, a stress responsive A/B barrel domain protein. **(b)** Raw electrical signals of poly(mA) and poly(A) detected from one poly(mA)-tailed poly(A) RNA isoform of Potri.010G150500. **(c)** Poly(A) and poly(A)^-^ RNA populations captured by poly(mA) tailing reflect the RNA content of cellular RNA samples. **(d)** Most of poly(A)^-^ RNAs captured by poly(mA) tailing had secondary alignments on *Populus* genome.

As expected, the majority of reads (over 72%) from control libraries were classified as “PASS”, proving that only poly(A) RNAs were targeted for sequencing in control samples (Fig. 3c). For each poly(mA)-tailed RNA library, we found that the amount of poly(A) and poly(A)^-^ RNA identified basically reflected its RNA composition. For example, for total RNA samples mainly consisting of poly(A)^-^ rRNAs, 87% of reads from the poly(mA)-tailed total RNA sample and over 72% of reads from the rRNA depleted total RNA sample, were classified as “NOREGION” without poly(A) tails (Fig. 3c). The ratio of poly(A)^-^ RNAs to poly(A) RNAs captured in the poly(mA)-tailed total RNA sample was 87% to 6%, close to the general projection for the total cellular RNA composition: 80-90% poly(A)^-^ rRNAs to 2-5% poly(A) mRNAs. For mRNA samples, RNAs were composed of mostly poly(A)-enriched RNAs and variable amounts of poly(A)^-^ RNAs from total RNA carryover during inefficient oligo(dT)-based poly(A) enrichment. Reasonably, from the two tailed mRNA libraries, we identified 47%-53% of reads were poly(A) RNAs, and at least 25% to 31% of reads were poly(A)^-^ RNAs. The overall results demonstrate the capability of poly(mA) tailing to capture the depth of RNA population in varying cellular contexts (Fig. 3c). Genome alignment of reads from each library further revealed that most poly(A)^-^ RNAs captured by poly(mA) tailing had multiple alignments to the *Populus* genome (Fig. 3d), confirming the repetitive nature of the majority of poly(A)^-^ RNAs captured in natural cellular RNA samples.

### Accessing organellar-origin RNA species on nanopore by poly(mA) tailing

In all poly(mA)-tailed libraries, we detected a significant number of reads mapped to the chloroplast genome of *Populus*; in contrast, many fewer reads from control libraries were aligned to the chloroplast genome (Fig. 4a). Of all the chloroplast RNAs sequenced, most were poly(A)^-^ RNAs (Fig. 4b). Indeed, further analysis of the read coverage of poly(mA)-tailed total RNAs across the chloroplast genome revealed the extensive transcription of poly(A)^-^ RNAs (Fig. 4c). Interestingly, we also observed that chloroplast-origin poly(A)^-^ RNA reads were overall longer than their polyadenylated isoforms (Fig. 4d).

**Figure 4.**
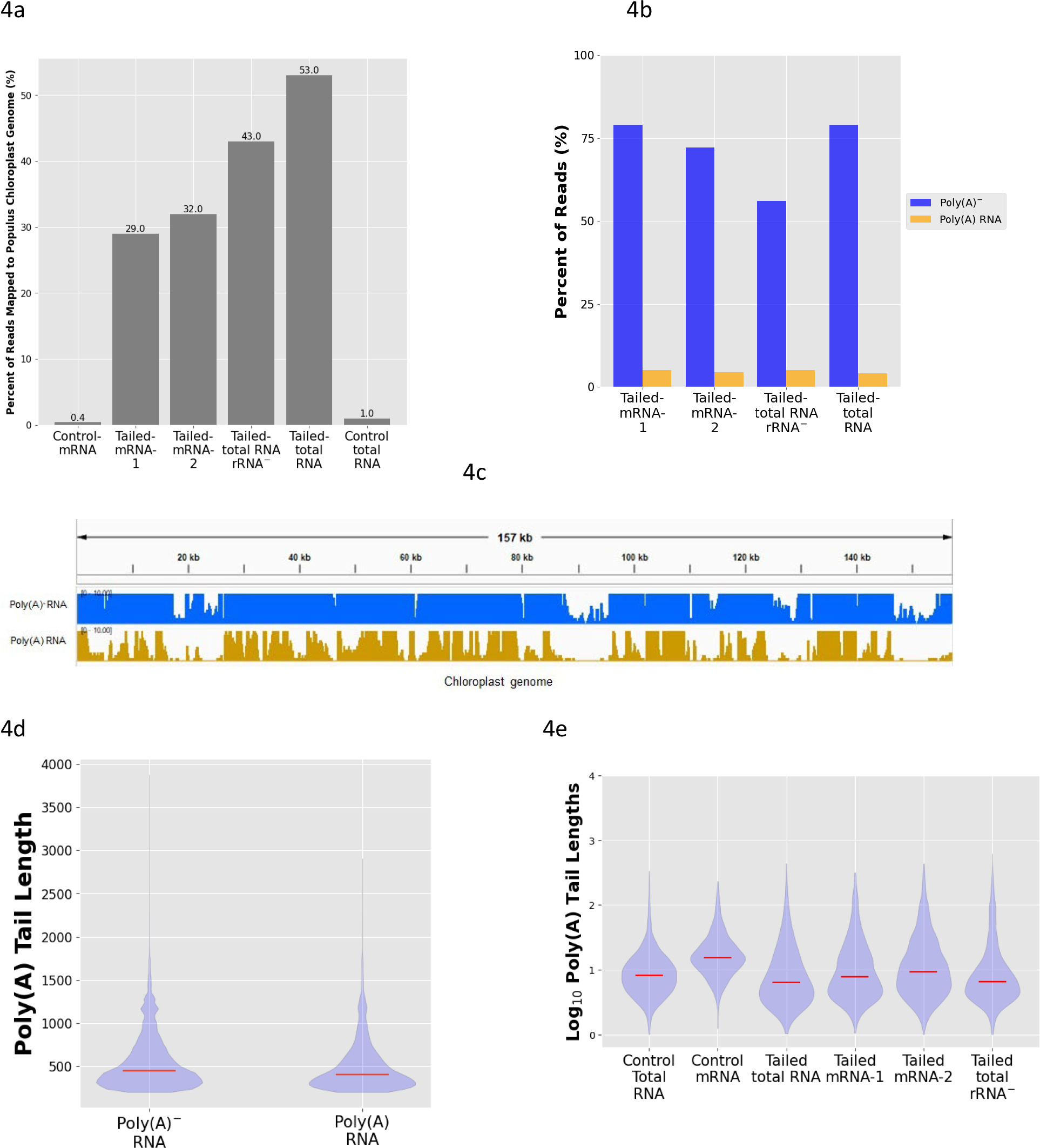
Chloroplast-origin RNAs were effectively captured by poly(mA) tailing. **(a)** Reads mapped to chloroplast genome were enriched in all poly(mA)-tailed RNA libraries. **(b)** Chloroplast-origin RNAs were predominantly poly(A)^-^ in poly(mA)-tailed RNA libraries. **(c)** Read coverage of poly(A)^-^ RNAs vs. poly(A) RNAs on *P. trichocarpa* chloroplast genome. All reads were derived from the poly(mA)-tailed total RNA (rRNA^-^) sample. **(d)** The violin plot shows the distribution of transcript length between chloroplast poly(A)^-^ and poly(A) RNA reads of poly(mA)-tailed total RNAs(rRNA^-^) library, with the mean transcript length indicated by the red line. (e) The violin plot shows distribution pattern of chloroplast poly(A) RNA tail length among control and poly(mA)-tailed libraries. The mean tail length is indicated by the red line.

Of the small number of chloroplast-origin poly(A) RNAs identified, we found that the median poly(A) tail length was less than 10 nt in most of the samples (Fig. 4e). The polyadenylation of the chloroplast RNAs, however, is complex. We identified some chloroplast RNAs contained poly(A) tail longer than 200 nt. The electrical signals of most of these long poly(A) tails on nanopore were also noisy and the polyadenylation patterns were quite distinct from those RNAs transcribed from the nuclear genome (Fig. 5). Some of the poly(A) signals appeared internal with a varying number of non-adenosine or modified nucleotides intermixed with few adenosines at 3’ end (Fig. 5b). Most of internal poly(A) RNAs were identified specifically in the poly(mA)-tailed total RNA library (Supplementary Table S2), and rarely detected in the mRNA control library, suggesting the complex chemical composition of 3’ end of such RNAs hinders their capture through the standard ONT mRNA sequencing approach.

**Figure 5.**
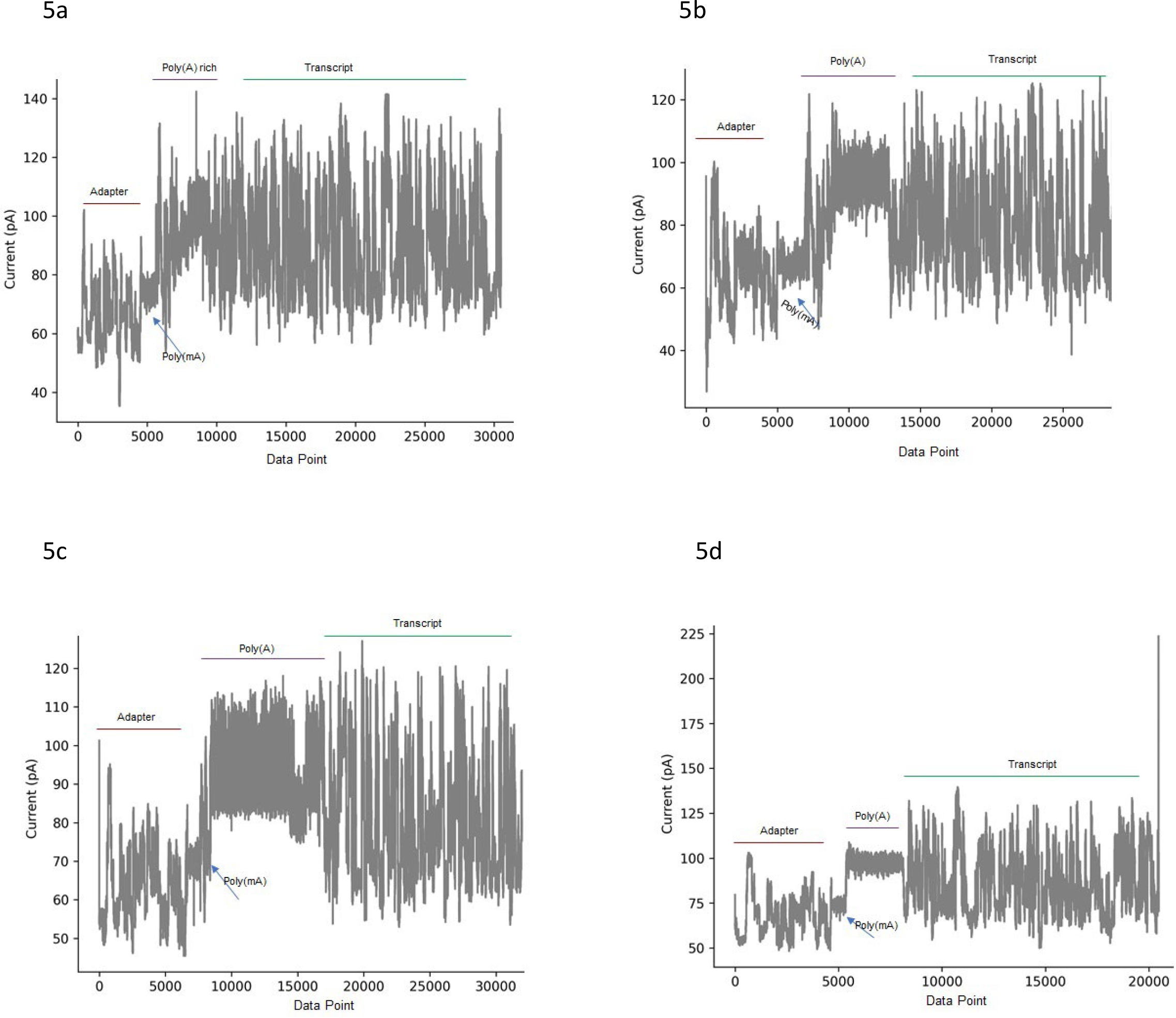
Nanopore ionic current traces of chloroplast-origin **(a-c)** vs. nuclear-origin poly(A) RNA **(d). (a)** A noisy poly(A)-rich signal was identified in one RNA isoform of chloroplast gene rbcL encoding ribulose bisphosphate carboxylase large chain precursor. **(b)** A defined length of internal poly(A) signal was detected in another RNA isoform of chloroplast gene rbcL. **(c)** A much wider current variation of poly(A) signal was observed with one RNA isoform of chloroplast gene psbA, photosystem II D1-reaction-center protein. **(d)** The classical poly(A) current traces was routinely identified in nuclear transcribed RNAs, such as this RNA isoform of rbcS, a nuclear gene Potri.004G100000 encoding ribulose bisphosphate carboxylase small chain 1A which targets chloroplast.

With its robust ability in capturing various types of chloroplast RNAs, poly(mA) tailing offers a necessary method for direct nanopore sequencing and characterizing such organellar RNAs for structural and functional analysis.

## DISCUSSION

Poly(mA) tailing was developed to address technical limitations in direct nanopore sequencing of total cellular RNAs. The advantages of this technical breakthrough include, (1) highly efficient poly(mA) tailing accesses total cellular RNAs, particularly poly(A)^-^ and organellar RNAs with complex 3’ end; (2) the distinct electrical signal of poly(mA) tail on nanopore retains native polyadenylation which are readily analyzable by common signal analysis tools; (3) the construction of poly(mA)-tailed RNA libraries is fully integrable with ONT direct RNA kit with uncompromised sequencing throughput.

One notable finding of our study is that YPAP efficiently incorporates 2′-OMeA, a modified adenosine with a methyl group added to the 2’ hydroxyl of the ribose moiety, into the 3’ end of RNA. 2’-O-methylation, a common nucleotide modification present in natural RNAs [24, 25], has not been used for in vitro 3’ end RNA tailing. Non-poly(A) tailing methods which generally use nucleotides with difference from adenosine in base structures, are not highly efficient in tailing poly(A)^-^ RNA, as has been reported [15] and also confirmed by our study (Supplemental Information, Supplementary Fig. S2a-d). The chemical structure of 2’-OMeA might underly the unique enzymatic activity of YPAP in adding poly(mA) tail to the 3’ end of RNA. As demonstrated in this study, despite its preference for poly(A)^-^ RNA, YPAP maintains processivity and distributivity in adding 2′-OMeA to both poly(A) and poly(A)^-^ RNA with a defined tail length by controlling the tailing reaction time (Fig. 1d). To note, because the poly(mA)-tailed RNA anneals to the poly(dT) adapter of the standard ONT direct RNA kit, native poly(A) RNAs are theoretically captured on nanopore even with inefficient poly(mA) tailing of poly(A) RNAs. Thus, integration of poly(mA) tailing with current ONT direct RNA sequencing technology not only captures poly(A)^-^ RNA with high efficiency, but also effectively retains poly(A) RNAs on nanopore, which sets poly(mA) tailing apart from other non-poly(A) tailing methods.

Similarly, likely due to its distinctive structure from other base-modified nucleotides, we found that of several polynucleotide tails tested, poly(mA) is the only one that generates distinct ionic current traces on nanopore that can be readily differentiated and segmented from the native poly(A) signal by nanopolish (Supplementary Fig. S3). Due to this unique feature, the native 3’ end polyadenylation status of poly(A) RNAs such as poly(A) tail length can be conveniently analyzed with current signal analysis tools. We demonstrated this utility through poly(mA) tailing of complex chloroplast-origin RNA species which are richly expressed in leaf cell but inefficiently captured through regular mRNA sequencing methods. Our analysis offered a direct peek at the pervasive transcription of poly(A)^-^ RNAs in chloroplast and the polyadenylation dynamics of chloroplast-origin RNA isoforms (Fig. 5). Polyadenylation regulates gene expression through modulating mRNA stability in all living organisms [26, 27]. There is a strong interest in understanding the coordinated gene expression between chloroplast and nuclear genomes [28]. Tracking the dynamics and heterogeneity of polyadenylation of organellar-origin RNAs could shed light on RNA metabolism and its role in regulating gene expression between these two genomes. We confirmed that poly(mA) tailing captures chloroplast RNAs as effectively as poly(A) tailing in various cellular contents (Supplementary Fig. S4), while retaining the complex nature of native poly(A) signal of individual RNA isoform, making it an essential tool for such applications.

Another potential limiting factor associated with 3’ custom tailing is the restricted sequencing throughput due to the long stretch of homopolymer tail inadvertently added to the 3’ end of RNA interfering with RNA threading through the nanopore [15]. Poly(mA) tailing, however, overcomes this issue easily. Our enzyme activity analyses have shown that the length of poly(mA) tail added by YPAP has been consistently controlled into a narrow range even with extended reaction times (Fig. 1d, Supplementary Fig. S2a-b), a strong contrast to poly(A) tailing and PUP mediated poly(I) or poly(U) tailing (Supplementary Fig. S2c, 2e, and 2g). Directly sequencing poly(mA)-tailed RNAs side-by-side with other custom tailed RNAs revealed the length of poly(mA)-tailed RNA strands sequenced on the device was confined to a much shorter range than those of poly(A) or poly(I) tailed RNA libraries (Supplementary Fig. S5). With the current libraries that we tested, poly(mA)-tailed RNAs outperformed other custom tailed RNAs with improved sequencing throughput (Supplementary Tables S3-4). The total number of reads obtained from the poly(mA)-tailed total RNA (rRNA^-^) sample on nanopore in this study was around one million reads, close to the typical number of reads from a standard ONT direct RNA kit per flow cell (Supplementary Table S5). Additionally, we found that although the poly(mA) tail anneals to the poly(dT) adapter, increasing the ligation time between them from 10 min to 15 min has been shown to improve the library yield. Thus, with the increased library input on the flow cell, the sequencing throughput of poly(mA) tailed RNA library is expected to be further improved.

Aside from capturing the total cellular RNA population on nanopore, we believe poly(mA) tailing will also have immediate benefits for other research fields, such as tracking the expression and regulation of nascent RNA transcription in cell, or any researches that require efficient 3’ end custom tailing to capture the newly synthesized, intermediate or chemically modified RNA species on nanopore for analysis.

## METHODS

### Template preparation and SP6 *in vitro* transcription of λRNA

DNA templates for in vitro transcription of λRNA were generated from a defined region of λDNA (NEB, N3011S) through PCR amplification. Template DNAs were amplified with a forward primer (5’-ATTTAGGTGACACTATAGAAGGTTCAGGGTTGTCGGACTTG -3’) containing a SP6 promoter region and either a reverse primer (5’-TGGCGAACAACAAGAAACTG -3’) terminating the targeted region for poly(A)^-^ λRNA, or a reverse primer (5’-TTT TTT TTT TTT TTT TTT TTT TTT TGGCGAACAACAAGAAACTG -3’) to add a 24 nt 3’ poly(A) tail to poly(A)^-^ λRNA for transcription of poly(A24) λRNA. PCR amplification was performed using Phusion™ Hot Start II DNA Polymerase (Thermo Fisher Scientific, F549S) in a reaction containing 1 x Phusion HF Buffer, 2 ng/μl λDNA, 0.2 mM dNTP mix, 0.5 μM forward primer, 0.5 μM reverse primer, and 0.02 U/μL of Phusion Hot Start II DNA Polymerase, with initial denaturation at 98 °C for 30s, 33 cycles of [98°C 5s, 58°C 10s, 72°C 15s], then 72°C for 10 min. The amplified and gel-verified PCR product was purified using AMPure XP beads (Beckman, A63880) following the manufacturer’s instructions and was then used for in vitro transcription.

In vitro transcription was carried out using HiScribe® SP6 RNA Synthesis Kit (NEB, E2070S) in a 25 μl reaction containing 1 x SP6 Reaction Buffer, 5 mM of each ATP, CTP, GTP and UTP, 1 μg of template DNA, and 2.5 μl SP6 RNA Polymerase Mix. The reaction was incubated at 37°C for 2 hours. The transcribed RNA was then separated on a 6% TBE-Urea (7M urea) polyacrylamide gel (Thermo Fisher Scientific, EC6865BOX) which was post stained with GelRed (Biotium, 41003). Transcribed RNA of the expected size was excised from the gel. Gel slices were mixed with RNA elution buffer (0.3 M NaOAc, pH 5.2, 0.2% SDS, 1 mM EDTA, 10 μg/mL proteinase K) at 4°C overnight. The eluted RNA was extracted with phenol and chloroform, and the eluted RNA was dissolved in RNase-free water for 3’ end custom tailing experiments.

### 3’ end polynucleotide tailing with YPAP

A 20-mer RNA oligo with FAM attached to the 5’ end (5’-/56-FAM/rGrCrUrArUrGrUrGrArGrArUrUrArArGrUrUrArU-3’) was custom-ordered from IDT. The standard reaction to add a polynucleotide tail of ATP or a modified analog to the 20-mer RNA oligo was set as follows: 36 pmol of RNA oligo added to a 20 μl reaction containing 1 x YPAP reaction buffer, 1 mM ATP or a modified ATP analog (Trilink, 2’-O-Methyladenosine-5’-Triphosphate, N-1015-10; 2-Aminoadenosine-5’-Triphosphate, N-1001-5; N6-Methyladenosine-5’-Triphosphate, N-1013-1; N1-Methyladenosine-5’-Triphosphate, N-1042), and 600 units of YPAP (Thermo Fisher Scientific, 74225Z25KU). The reaction was incubated at 37°C for 5 min. Reactions were terminated with 5 mM EDTA, and mixed with an equal volume of 2 x RNA loading dye (NEB, B0363S). 25 μl of the mixture was loaded on a 10% TBE-Urea gel (Bio-Rad, 4566033), together with the low range ssRNA ladder (NEB, 0364S).

To add polynucleotide tail of ATP or modified analog to λRNA, the reaction was set up similarly as with 20-mer RNA oligo except that 1.9 pmol of RNA was used per 20 μl reaction, and half of the reactions were incubated at 37°C for 5 min and another half for 30 min. The reactions were stopped by adding 5 mM EDTA and mixed with equal volume of 2 x RNA loading dye. 25 μl mixture was heated at 80°C for 3 min, and then loaded on 6% TBE-Urea gel alongside the ssRNA ladder.

### 3’ end polynucleotide tailing of 20-mer RNA oligo with EPAP

To test the ability of EPAP in adding polynucleotide tail of ATP or modified ATP analog to the RNA oligo, 36 pmol RNA oligo were added to a 20 μl reaction containing 1 x EPAP reaction buffer, 1 mM ATP or modified analog, and 5 units of EPAP (NEB, M0276L). The reaction was incubated at 37°C for 5 min, stopped by adding 5 mM EDTA, and mixed with 1 x RNA loading buffer. 25 μl of the mixture was loaded on a 10% TBE-Urea gel.

### Poly(mA) tailing of λRNA mixture for a time-series experiment

The mixture of poly(A)^-^ λRNA and its poly(A24) form (at a ratio of 2 to 1), was used as the substrate by YPAP in adding poly(mA) tails for a time-series experiment. 4.61 pmol of λRNA mixture was mixed with 1 x YPAP reaction buffer, 1 mM 2′-OMeA, and 1800 units of YPAP in a 30 μl volume. The 30 μl reaction mixture was then split into six 5 μl reactions, and each reaction was incubated at 37°C for the following specified time point: 0 min (control), 5 min, 30 min, 60 min, 90 min, and 120 min. At each time point, the reaction was terminated by mixing with 5mM EDTA and 1 x RNA loading dye. All six reactions were loaded on a 6% TBE-Urea gel. The gel was post-stained with GelRed, and visualized with a UVP GelDoc-It imaging system.

The same RNA substrate mixture was also used for EPAP using 2′-OMeA. The poly(mA) tailing reaction was performed in the same way as the YPAP time-series experiment, except that the 30 μl reaction mixture was in 1 x EPAP reaction buffer and contained 15 units of EPAP. Similarly, the 30 μl reaction mixture was split into six 5 μl reactions, and each reaction was incubated at 37°C for the specified time (0 min, 5 min, 30 min, 60 min, 90 min, or 120 min).

### Plant RNA extraction, mRNA purification and rRNA depletion

Total RNA from *P. trichocarpa* leaf tissue was extracted through CTAB method [29] and cleaned up with RNAClean XP beads (Beckman Coulter, A63987) after DNase I treatment (Thermo Fisher Scientific, EN0521). Poly(A) mRNA was selected using Oligo d(T)25 Magnetic Beads (NEB, S1419S) following the manufacturer’s instructions. Ribosomal RNAs were depleted from DNase I treated total RNAs by employing the riboPOOL kit for plants (siTOOLs Biotech, dp-K012-000031) following the manufacturer’s instructions. The recovered rRNA depleted total RNA was further cleaned up and concentrated with the RNA Clean & Concentrator kit (Zymo, R1016).

### Poly(mA) tailing of cellular RNA sample with yeast poly(A) Polymerase

To add poly(mA) tails to cellular RNAs, 300 to 500 ng of enriched poly(A) mRNA or rRNA-depleted total RNA or 1 to 1.5 μg of DNase I treated total RNA, was first heated to 65°C for 5 min to denature any secondary structure of the RNAs and used in a 20 μl reaction volume containing 1 x YPAP reaction buffer, 1 mM 2′-OMeA, and 600 units of YPAP. The reaction was incubated at 37°C for 60 min and terminated with 5mM EDTA. The tailing reaction was cleaned up with RNAClean XP beads, and the tailed RNA was then eluted in water for further use. Additionally, to make sure that 2′-OMeA was completely removed from tailed RNA samples, the eluted, tailed RNA was further desalted by running on Performa spin columns (EdgeBio, 73328) following the manufacturer’s instructions.

### Poly(mA) and poly(A) tailing of λRNA for direct RNA library construction and nanopore sequencing

Three types of 3’ end custom tailed λRNA were prepared for direct RNA library construction.

#### Poly(mA) tailed poly(A)^-^ λRNA

Approximately 400 ng of poly(A)^-^ λRNA were tailed with poly(mA) by YPAP in a 20 μl standard reaction as described earlier. The reaction was incubated at 37°C for 60 min. Tailed RNA was cleaned up and desalted as described in cellular RNA tailing experiments.

#### Poly(A) tailed λRNA (poly(A) λRNA), and poly(mA) tailed poly(A) λRNA

Approximately 500 ng of poly(A)^-^ λRNA were tailed using ATP by EPAP in a 20 μl reaction containing 1 x EPAP reaction buffer, 1 mM ATP, and 15 units of EPAP at 37°C for 2 min. The poly(A) tailed RNA was then cleaned up with RNAClean XP beads and eluted in 20 μl of water. 10 μl of the eluted poly(A) tailed RNA was used for direct RNA library construction, and the rest was used for poly(mA) tailing with YPAP through standard poly(mA) tailing procedure.

The tailed λRNAs prepared above were employed for direct RNA library construction using the ONT SQK-RNA002 kit. The libraries were made without the optional RT step, and each library was sequenced on a MinION sequencer with a Flongle Flow Cell (R9.4.1) (FLO-FLG001), running the standard MinKNOW software with the base-calling tab set to OFF.

### Direct cellular RNA library construction for sequencing on nanopore

#### ONT standard RNA library

1 μg of DNase I treated total cellular RNA or 500 ng of enriched mRNA was used for ONT standard RNA library construction.

#### Poly(mA) tailed RNA library

500 to 750 ng of poly(mA) tailed total RNA, 300 to 400 ng of poly(mA) tailed mRNA, and 100 to 300 ng of poly(mA) tailed total cellular RNA with rRNA depleted were prepared for direct RNA library construction.

All libraries were constructed using the ONT SQK-RNA002 kit following ONT ‘s full protocol. For the reverse transcription step, we used Maxima H Minus Reverse Transcriptase (Thermo Fisher Scientific, EP0752). Libraries were sequenced on the MinION sequencer with a ONT R9.4.1 flow cell (FLO-MIN106D) running the standard MinKNOW software with the base-calling tab set to OFF. For libraries run on the same flow cell, the ONT flow cell wash kit (EXP-WSH004) was used between runs to remove the previous library from the flow cell.

### RNA reads base calling and mapping to reference genome

Raw fast5 files generated from the MinION sequencer were base called with Guppy Basecalling Software, version 6.4.6+ae70e8f, provided by Oxford Nanopore Technologies, with the configuration file “rna_r9.4.1_70bps_hac” [30]. The base called λRNA reads in fastq format were then mapped to the J02459.1 Escherichia phage Lambda reference genome through minimap2 version 2.17-r941 [31] with the parameter: -ax map-ont -k14. RNA reads from *Populus* RNA libraries were mapped to the *Populus* reference genome v3.0 with the parameters: -a -x splice -k14 -uf, or to the *Populus* chloroplast genome EF489041.1 with the parameters: -ax map-ont -k14. For downstream expression analysis, the aligned sam file was further filtered with samtools [32]: samtools view -h -F 2324, to remove unmapped, secondary and supplementary reads.

### Nanopolish classification and poly(A) tail length estimation

Nanopolish version 0.14.0 was used to classify RNA reads into poly(A) and poly(A)^-^ transcripts. First fast5 reads and base-called fastq files were indexed through nanopolish index version 0.14.0 with the default parameters, then nanopolish polya was called to classify reads into “PASS”, “NOREGION”, or “ADAPTER” and to estimate the poly(A) tail length of “PASS” reads containing poly(A) tails.

### Nanopore raw signal data manipulation and visualization

Raw signal of nanopore direct RNA-seq reads were routinely manipulated with ont_fast5_api (https://github.com/nanoporetech/ont_fast5_api) and SquiggleKit [33]. SquigglePlot was used to plot the raw signal data for visualization.

## Supporting information

Supplementary Figures, Tables, Methods and Other Information

## Data availability

Fast5 and fastq files generated in this study have been submitted to NCBI Sequence Read Archive (SRA) under the BioProject accession number PRJNA987605 and PRJNA1023552.

https://dataview.ncbi.nlm.nih.gov/object/PRJNA987605?reviewer=dqba2hrlhfkvcljm4mrv95nt0o

https://dataview.ncbi.nlm.nih.gov/object/PRJNA1023552?reviewer=3076ljqqgikscos9t4rvoi8vco

