## Supplementary Figures, Tables, Methods and Other Information for "Efficient 3ʹ-end tailing of RNA with modified adenosine for nanopore direct total RNA sequencing"

**Supplementary Table S1.** Overview of sequencing run and mapping information for natural cellular libraries

| Library | Run time (hr) | Reads generated | Base called pass reads | Reads mapped to *Populus* genome |
|---|---|---|---|---|
| Control-mRNA | 24 | 1,610,000 | 1,509,527 | 1,469,115 |
| Tailed-mRNA-1 | 11.74 | 436,340 | 407,755 | 388,226 |
| Tailed-mRNA-2 | 11.94 | 410,560 | 365,974 | 338,158 |
| Tailed-total rRNA$^-$ | 40.16 | 960,010 | 824,174 | 682,255 |
| Tailed-total RNA | 24 | 119,590 | 101,207 | 91,098 |
| Control-total | 42 | 2,550,000* | 2,548,850* | 1,135,214 |

*Note: Calibrant Strand (RCS) RNA from ONT kit was over-sequenced in this library. We didn't include RCS in other libraries.

**Supplementary Table S2**. Varying number of organellar-origin poly(A) RNAs mapped to several chloroplast-related gene loci in control and poly(mA)-tailed RNA libraries.

| Chloroplast-related gene loci | Tailed-total RNA (rRNA$^-$) | Tailed-mRNA-1 | Control-mRNA | Control-total RNA |
|---|---|---|---|---|
| Potri.011G075000 | 25 | 12 | 1 | 21 |
| Potri.013G138000 | 6 | 2 | 0 | 3 |
| Potri.013G143300 | 7 | 4 | 6 | 3 |
| Potri.013G137700 | 5 | 8 | 0 | 7 |
| Potri.013G138900 | 5 | 2 | 3 | 2 |
| Potri.013G142800 | 8 | 1 | 0 | 0 |
| Potri.013G138800 | 27 | 0 | 1 | 1 |
| Potri.013G137900 | 10 | 5 | 0 | 1 |
| Potri.013G142100 | 273 | 3 | 5 | 14 |
| Potri.013G139000 | 3 | 4 | 0 | 2 |
| Potri.005G154600 | 1 | 0 | 0 | 0 |
| Potri.005G154700 | 20 | 7 | 1 | 7 |
| Potri.012G062600 | 1147* | 3 | 0 | 1 |
| Potri.005G154900 | 12 | 5 | 5 | 11 |
| Potri.011G074200 | 76 | 142 | 19 | 98 |

*Note: significant number of internal poly(A) RNAs mapped to this gene locus.

**Supplementary Table S3.** Run summary of different types of RNA libraries on flow cell one.

| Run order | Library type | Amount of library loaded (ng) | Run time (hr) | Data produced | Reads generated | Run performance rank |
|---|---|---|---|---|---|---|
| 1 | Nanopore standard direct mRNA library | 130 | 24 | 62.49 GB | 1.61 M | 1 |
| 2 | Poly(mA)-tailed-mRNA library-1 | 40 | 11.74 | 15.22 GB | 436.34 K | 2 |
| 3 | Poly(I)-tailed-mRNA library | 34 | 9.38 | 1.56 GB | 42.75 K | 4 |
| 4 | Poly(mA)-tailed-mRNA library-2 | 36 | 11.46 | 5.04 GB | 147.94 K | 3 |

**Supplementary Table S4.** Run summary of different types of RNA libraries on flow cell two.

| Run order | Library type | Amount of library loaded (ng) | Run time (hr) | Data produced | Reads generated | Run performance rank |
|---|---|---|---|---|---|---|
| 1 | Poly(mA)-tailed-mRNA library | 30 | 11.94 | 12.11 GB | 410.56 K | 1 |
| 2 | Poly(A)-tailed-mRNA library | 33 | 11.34 | 2.78 GB | 89.61 K | 3 |
| 3 | Poly(mA)-tailed-total RNA library | 51 | 17.65 | 8.01 GB | 218.22 K | 2 |

**Supplementary Table S5.** Sequencing poly(mA)-tailed total RNAs on flow cell three.

| Run order | Library type | Amount of library loaded (ng) | Run time (hr) | Data produced | Reads generated |
|---|---|---|---|---|---|
| 1 | Poly(mA)-tailed-total RNA ($rRNA^-$) library | 38 | 40 | 22.61 GB | 906.01 k |
| 2 | Poly(mA)-tailed-total RNA library | 37 | 47 | 10.86 GB | 266.43 k |

**Supplementary Table S6.** Capturing chloroplast RNAs from same RNA sample through poly(mA) tailing vs. other tailing methods.

| Dataset | Reads generated | Reads mapped to *Populus* chloroplast genome |
|---|---|---|
| Poly(A)-tailed total | 138450 | 103926 |
| Poly(mA)-tailed total-1 | 268610 | 218761 |
| Poly(mA)-tailed total-2 | 168490 | 132873 |
| Poly(I)-tailed total | 283510 | 181842 |

### Supplementary Figures

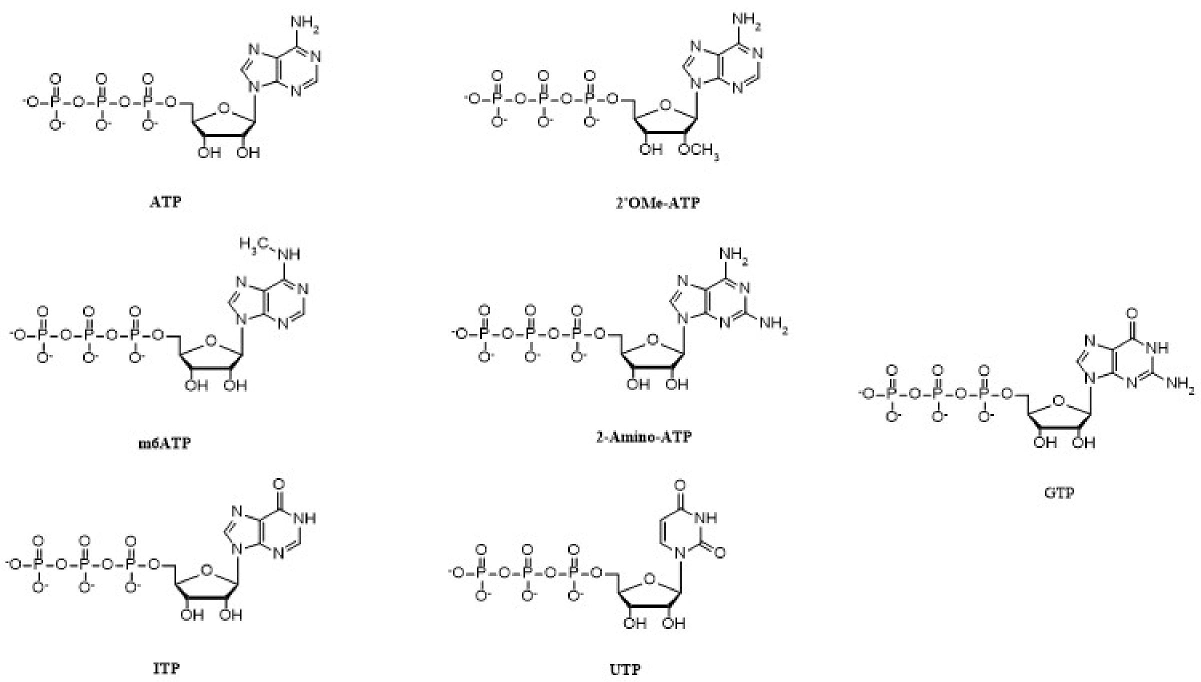


**Supplementary Figure S1.** Chemical structures of ATP, modified ATP analogs, and other nucleotides used for 3’ end RNA tailing in this study.

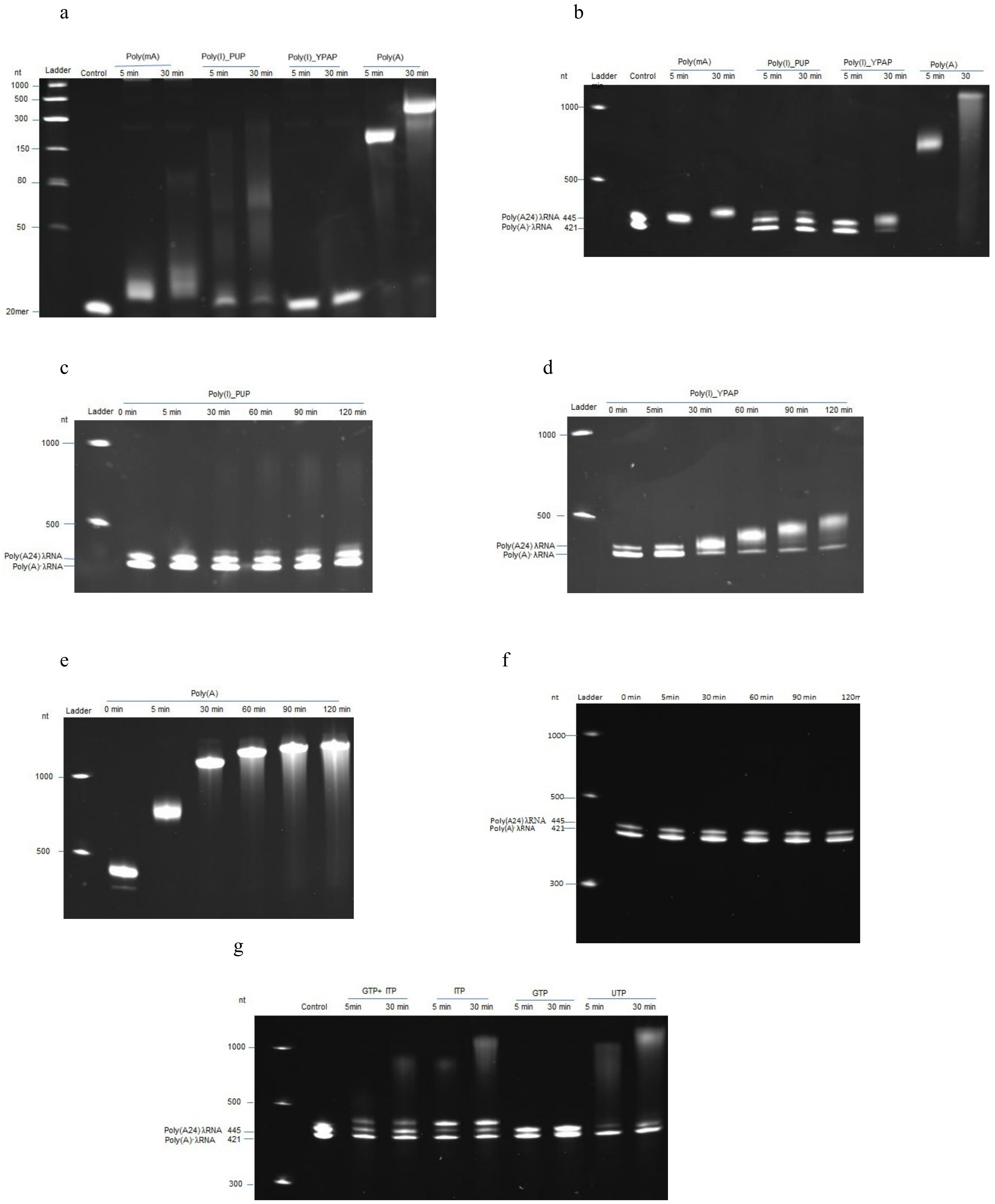

a
nt
Ladder
Control
Poly(mA)
5 min
30 min
Poly(I)_PUP
5 min
30 min
Poly(I)_YPAP
5 min
30 min
Poly(A)
5 min
30 min
1000
500
300
150
80
50
20mer
b
nt
Ladder
Control
Poly(mA)
5 min
30 min
Poly(I)_PUP
5 min
30 min
Poly(I)_YPAP
5 min
30 min
Poly(A)
5 min
30
1000
500
Poly(A24) λRNA 445
Poly(A)⁻ λRNA 421
c
Poly(I)_PUP
nt
Ladder
0 min
5 min
30 min
60 min
90 min
120 min
1000
500
Poly(A24) λRNA
Poly(A)⁻ λRNA
d
Poly(I)_YPAP
Ladder
0 min
5min
30 min
60 min
90 min
120 min
1000
500
Poly(A24) λRNA
Poly(A)⁻ λRNA
e
Poly(A)
Ladder
0 min
5 min
30 min
60 min
90 min
120 min
nt
1000
500
f
nt
Ladder
0 min
5min
30 min
60 min
90 min
1000
500
Poly(A24) λRNA 445
Poly(A)⁻ λRNA 421
300
g
nt
Control
GTP+ ITP
5min
30 min
ITP
5 min
30 min
GTP
5 min
30 min
UTP
5 min
30 min
1000
500
Poly(A24) λRNA 445
Poly(A)⁻ λRNA 421
300

**Supplementary Figure S2.** Assessing the efficiency of other tailing methods vs. poly(mA) tailing. **(a)** The enzyme activities of YPAP in adding a poly(mA) tail or poly(I) tail (Poly(I)_YPAP), and PUP in adding a poly(I) tail (Poly(I)_PUP), to a 20-mer non-poly(A) RNA oligo, were compared side-by side with the EPAP mediated poly(A) tailing. In contrast to its high efficiency in adding poly(mA) tail, YPAP showed extremely low efficiency in adding a poly(I) tail to the 20-mer substrate. **(b)** The efficiency of YPAP mediated poly(I) and PUP mediated poly(I) tailing were evaluated using a substrate mixture of equal amounts of poly(A)$^-$ λRNA and poly(A) λRNA, alongside poly(mA) tailing by YPAP and poly(A) tailing by EPAP, at 5 min and 30 min of reaction. The difference in tailing efficiency was observed from the highly efficient poly(mA) tailing and poly(A) tailing to low efficient poly(I) tailing. **(c)** A time-series of PUP mediated poly(I) tailing was performed on a substrate mixture of poly(A)$^-$ λRNA and poly(A) λRNA (2:1). A large amount of uncatalyzed poly(A)$^-$ RNA substrates were identified even with a 2-hour reaction. **(d)** A time-series of poly(I) tailing by YPAP was performed on a substrate mixture of poly(A)$^-$ λRNA and poly(A) λRNA (2:1), with significant amount of poly(A)$^-$ λRNA substrate remaining uncatalyzed over a 2-hour reaction. **(e)** A time-series of EPAP mediated poly(A) tailing of a substrate mixture of poly(A)$^-$ λRNA and poly(A) λRNA (2:1), demonstrated the high reaction rate of EPAP in adding poly(A) tail to the 3' end of RNA. The intensity and band shifting pattern of the substrate mixture at 0 min indicated that EPAP mediated poly(A) tailing was highly reactive even at the reaction preparation step, and long stretches of high-molecular-weight tailed substrates were observed along the tailing reaction progressing at 37°C. These intrinsic features of poly(A) tailing are exactly opposite to the slow reaction rate and tightly controlled poly(mA) adding activity (as demonstrated in Fig. 1d) **(f)** The inability of EPAP in adding 2'-OMeA to a substrate mixture of poly(A)$^-$ λRNA and poly(A) λRNA (2:1) was demonstrated through a time-series of reactions. **(g)** The nucleotide-adding activity of PUP was investigated with various nucleotides (ITP, GTP and UTP) and a combination of GTP and ITP, using a substrate mixture of poly(A)$^-$ λRNA and poly(A) λRNA (1:1) at 5 min and 30 min reaction time. Across nucleotides that we tested, PUP displayed low efficiency in catalyzing poly(A)$^-$ RNA while easily synthesizing long tails onto a fraction of substrates over the 30 min reaction.

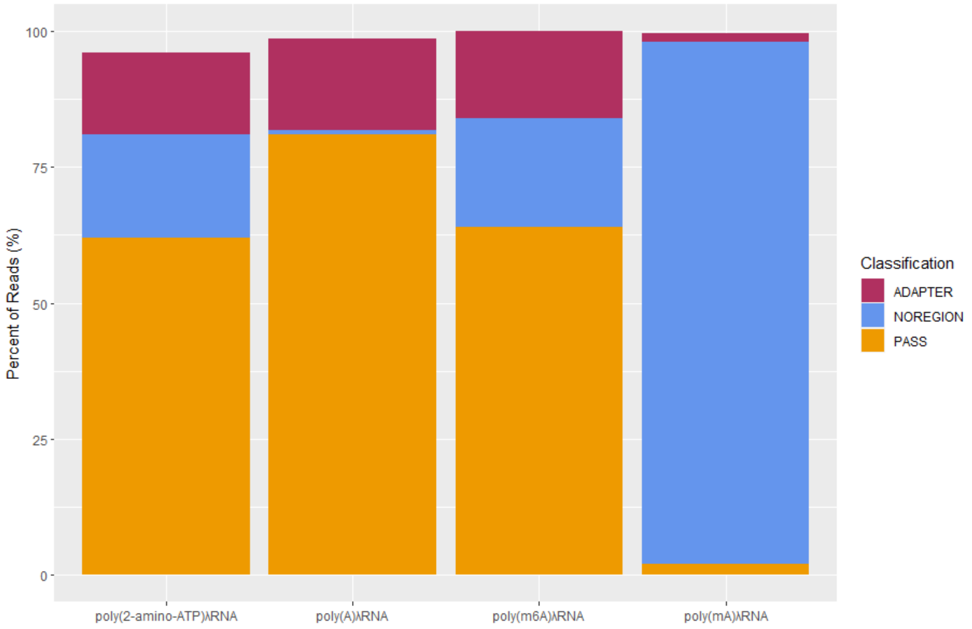


**Supplementary Figure S3.** Summary of nanopolish-polya analyses from nanopore direct RNA sequencing of four custom tailed λRNA libraries including 2-amino-ATP tailed (poly(2-amino-ATP)), m6A tailed (poly(m6A)), poly(A) tailed, and poly(mA) tailed λRNA. An overwhelming number of poly(mA) tailed RNA reads (>90%) were assigned into "NOREGION" through nanopolish segmentation analysis, revealing that the signal of poly(mA) tail is distinct enough from the signal of poly(A) tail; while over 60% of m6A tailed reads or 2-amino-ATP tailed reads were classified into the "PASS" category, suggesting the electric signal of polynucleotide of m6A or 2-amino-ATP was not distinct enough to be distinguished from the classic signal of poly(A) tail by the current nanopolish algorithm.

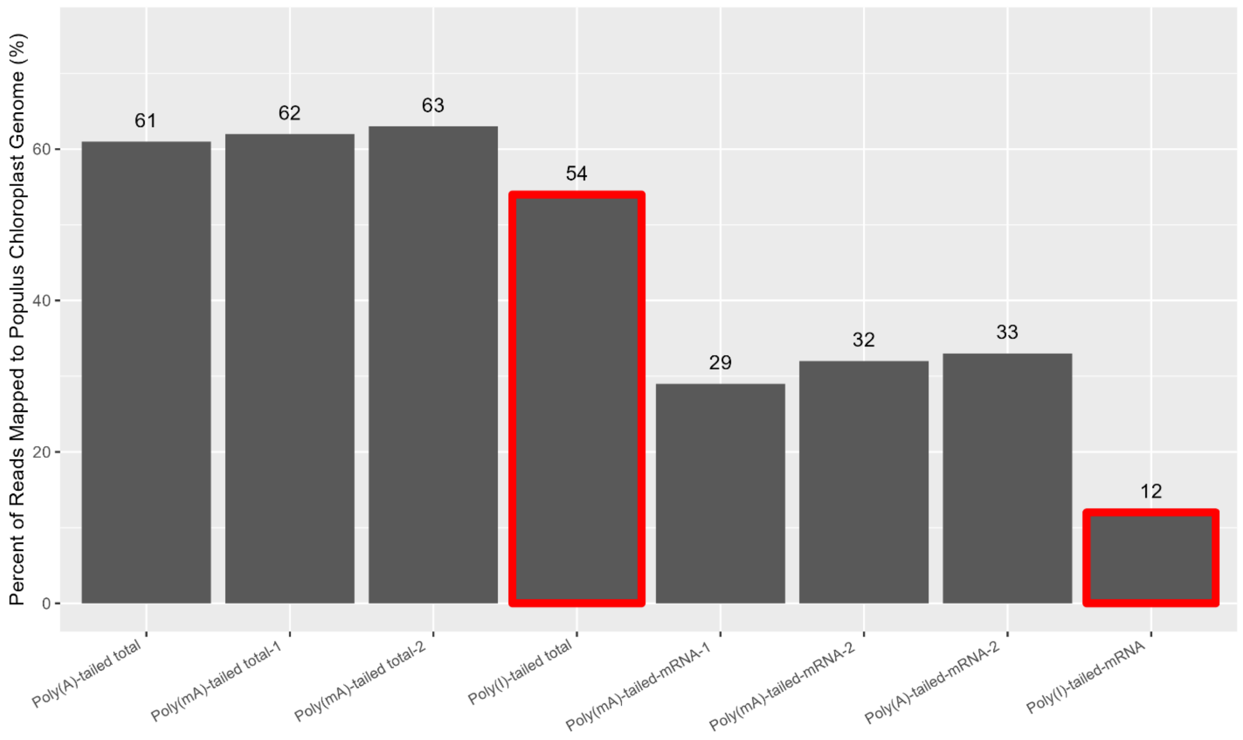


**Supplementary Figure S4.** The effectiveness of poly(mA) tailing in capturing chloroplast RNAs through nanopore direct RNA sequencing was assessed alongside poly(A) tailing and PUP mediated poly(I) tailing, from two types of cellular samples: total cellular RNA (datasets: Poly(mA)-tailed total, Poly(A)-tailed total and Poly(I)-tailed total. Library information was summarized in Supplementary Table S6.) and oligo(dT)-based poly(A)-enriched mRNA (datasets: Poly(mA)-tailed-mRNA, Poly(A)-tailed-mRNA, and Poly(I)-tailed-mRNA. Library information can be found in Supplementary Table S3 and S4.). By mapping the direct RNA sequencing reads from each library dataset to *Populus* chloroplast genome, we found that across cellular types, at the current sequencing depth assayed, poly(mA) tailing outperformed the poly(I) tailing (outlined in red line), particularly in the enriched mRNA content, and captured chloroplast RNAs as effectively as the poly(A) tailing.

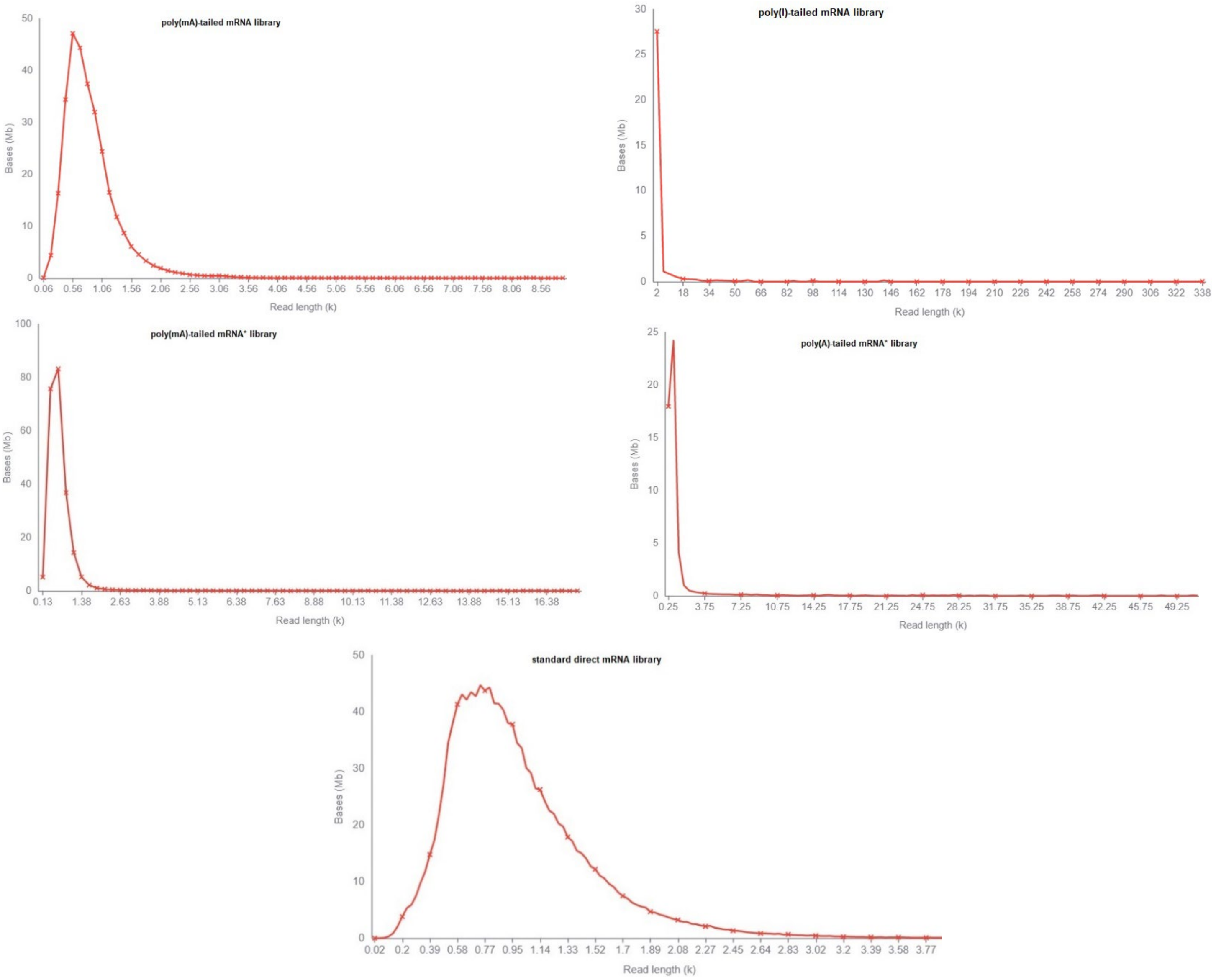


**Supplementary Figure S5.** The length graph was estimated by MinKNOW, the software that drives the nanopore sequencer, while the RNA strand threading through the nanopore at a steady rate controlled by the motor protein, showing the total number of bases (y-axis) vs. the read length (x-axis) after removing the 1% of outlier strands. The length graph of each library represents the raw read length distribution of sequenced RNAs on the sequencer. A length graph of a standard ONT direct mRNA library was included as a reference. For direct comparison, we generated one poly(mA)-tailed RNA library from the same enriched mRNA sample used for poly(I)-tailed RNA library, and another poly(mA)-tailed RNA library (labeled with an asterisk) was constructed from a different mRNA source (enriched mRNA samples was pretreated with DMSO at 65°C for 60 min.) which was used for a poly(A)-tailed library construction. In each case examined, the poly(mA) tailed RNA library displayed overall narrow and short length distribution, while the range of poly(I) or poly(A) tailed RNAs was quite extended (poly(I)-tailed RNA length range reaching over 300 kb on the x-axis was identified).

### Supplementary Methods and Additional Information

#### 3' end poly(I) tailing of a 20-mer RNA oligo

To test the efficiency of YPAP or PUP in synthesizing a poly(I) tail to the 20-mer RNA oligo, 36 pmol RNA oligo were added to a 20 μl reaction containing either 1 x YPAP reaction buffer or 1 x NEB reaction buffer 2, 1 mM ITP (Sigma, I0879), and 600 units of YPAP or 4 units of PUP (NEB, M0337S) in a 20 μl volume. Then the reaction mixture was split into two 10 ul reaction and incubated for 5 min and 30 min respectively. The reactions were stopped by mixing 5 mM EDTA and 1 x RNA loading buffer, and was loaded on a 10% TBE-Urea gel.

#### 3' end polynucleotide tailing of λRNA mixtures for time-series experiments

Time-series tailing reactions were performed to investigate the efficiency of YPAP or PUP mediated poly(I) tailing in tailing $poly(A)^-$ and poly(A) RNA using a mixture of $poly(A)^-$ λRNA and poly(A24) form (at a ratio of 2 to 1) as the substrate.

_Poly(I) tailing_: 4.61 pmol of λRNA mixture was mixed with 1 x YPAP reaction buffer or 1 x NEB reaction buffer 2, 1 mM ITP, and 1800 units of YPAP or 6 unites of PUP in a 30 μl volume. The 30 μl reaction mixture was then split into six 5 μl reactions, and each reaction was incubated at 37°C for the following specified time point: 0 min, 5 min, 30 min, 60 min, 90 min, and 120 min. At each time point, the reaction was terminated by mixing with 5mM EDTA and 1 x RNA loading dye. All six reactions were loaded on a 6% TBE-Urea gel. The gel was post-stained with GelRed, and visualized with a UVP GelDoc-It imaging system.

_Poly(A) tailing_: The same λRNA substrate mixture used for poly(I) tailing was also used to evaluate the EPAP mediated poly(A) tailing over a time series. The tailing reactions and gel analysis were performed similarly as poly(I) tailing except that 1 x EPAP buffer was used with 1mM ATP and 15 unites of EPAP.

#### Poly(U) polymerase catalyzing a substrate mixture of $poly(A)^-$ and ploy(A) RNA using various nulceotides

About 250 ng of RNA substrate mixture of poly(A)$^-$ λRNA and its poly(A24) form (at a ratio of 1 to 1), was used for PUP tailing experiment. The RNA substrate was mixed with 1 x NEB buffer 2, 1mM ITP or GTP (Thermo Fisher Scientific, R0461), UTP (NEB, M0337S), or the mixture of GTP and ITP (0.5 mM of each ITP and GTP), and 4 units of PUP in a 20 μl volume. The reaction mixture of each tested nucleotide or nucleotide mixture was then split into two 10 ul reaction and incubated for 5 min and 30 min respectively. The reactions were stopped with 5mM EDTA, and mixed with 10 ul 2 x RNA dye and loaded onto a 6% TBE-Urea gel.

**Poly(A)-tailed cellular RNA library construction**

300 to 400 ng of oligo-(dT) enriched mRNA, or 300 ng of enriched mRNA pretreated with DMSO at 65°C for 60 min, or 2 μg of total RNA or 500 ng of rRNA depleted total RNA, were tailed with poly(A) by EPAP in a 20 μl of reaction containing 1 x EPAP reaction buffer, 0.5 mM ATP, and 5 units of EPAP. The reaction was incubated at 37°C for 2 min following the recommended poly(A) tailing protocol by ONT (https://community.nanoporetech.com/extraction_methods/3-poly-rna-ecoli-pap). The tailing reaction was stopped by adding 5mM EDTA and purified with RNAClean XP beads. The purified poly(A)-tailed RNA was eluted in 11 μl water and ready for direct RNA library construction. The poly(A) tailed RNA library was constructed using the ONT SQK-RNA002 kit following the manufacture's full instructions.

**Poly(I)-tailed cellular RNA library preparation**

450 ng of oligo-(dT) enriched mRNA, or 2 μg of total RNA or 500 ng of rRNA depleted total RNA, were tailed respectively with poly(I) by PUP in a 20 μl reaction containing 1 x buffer 2, 0.5 mM ITP, and 4 units PUP. The reaction was incubated at 37°C for 30 min. The poly(I)-tailed RNA was purified through RNAClean XP beads and eluted in 11 μl water. For direct poly(I)-tailed RNA library construction, ONT SQK-RNA002 kit was used with a custom oligo(dC) adapter replacing the RTA adapter. The oligo(dC) adapter was assembled through annealing equimolar concentrations of the top oligo (5'-/5PHOS/GGCTTCTTCTTGCTCTTAGGTAGTAGGTTC-3') and the bottom oligo (5'-GAGGCGAGCGGTCAATTTTCCTAAGAGCAAGAAGAAGCCCCCCCCCCCC-3'), following ONT's instructions. The library was then constructed following ONT's instructions including the optional RT step.

**Mapping nanopore direct RNA sequencing data to *Populus* chloroplast genome**

Fast5 files of each custom tailed RNA libraries were generated from the MinION sequencer and was base called with Guppy. Base called poly(I) or poly(A) tailed reads were mapped through minimap2 to the *Populus* chloroplast genome EF489041.1 with the parameters: -ax map-ont -k14. The sam files produced from each library dataset were used to obtain the chloroplast genome mapping stats by running samtools flagstat.

**Assessing the sequencing throughput of poly(mA)-tailed RNA libraries on nanopore**

Low sequencing throughput has been a limiting factor for 3' end custom tailing-based direct nanopore RNA sequencing approaches. We performed the following experiments to examine the sequencing throughput of poly(mA)-tailed RNAs on nanopore. To avoid batch effects of flow cells, we decided to compare the run performance of poly(mA)-tailed RNAs with other types of RNA libraries on same R9.4.1 flow cell. As shown in Table S3, we evaluated one group of libraries including one standard ONT direct RNA library, two poly(mA)-tailed RNA libraries and one poly(I)-tailed RNA library on flow cell one (Tables S3). As shown in Table S4, we assessed another group of libraries including two poly(mA)-tailed

RNA libraries and one poly(A)-tailed RNA library on flow cell two. To run the libraries of each group continually on same flow cell, we loaded one library first and run for a period of time, then we performed flow cell wash before loading the next library for another run. This procedure was repeated in the order specified in Table S3 and S4 for each flow cell.

We then ranked the run performance of each library mainly based on the sequencing throughput (the number of reads generated for each library in the defined run time). When interpreting the run performance of each library, we also considered the fact that the quality of flow cell is going down gradually with the run time. Taken all data together from Tables S3 and S4, we concluded that poly(mA)-tailed RNAs perform much better than poly(I)-tailed RNAs and poly(A)-tailed RNAs on nanopore.

Lastly, we loaded one poly(mA)-tailed rRNA-depleted total RNA library and one poly(mA)-tailed total RNA library on a R9.4.1 flow cell (flow cell three) for a full examination of the sequencing throughput of poly(mA)-tailed RNAs. We obtained ~1.1 million raw reads from the flow cell three (Supplementary Table S5).
